# Soil collected in the Great Smoky Mountains National Park yielded a novel *Listeria* species, *L. swaminathanii*, effectively expanding the *sensu stricto* clade to ten species

**DOI:** 10.1101/2021.12.21.473762

**Authors:** Catharine R. Carlin, Jingqiu Liao, Lauren K. Hudson, Tracey L. Peters, Thomas G. Denes, Renato Orsi, Xiaodong Guo, Martin Wiedmann

**Affiliations:** Department of Food Science, Cornell University, Ithaca, NY 14853, USA; Department of Microbiology, Cornell University, Ithaca, NY 14853, USA; Department of Systems Biology, Columbia University, New York, NY 10032, USA; Department of Food Science, University of Tennessee, Knoxville, TN 37996, USA

**Author notes:** **Correspondence:** Martin Wiedmann. **Repositories:** The GenBank/EMBL/DDBJ accession numbers for the 16S rRNA and draft genome sequences for the type strain, FSL L7-0020^T^ are MT117895 and JAATOD000000000, respectively.

**Keywords:** *Listeria sensu stricto*, novel species, average nucleotide identity, *in silico* DNA-DNA Hybridization, US National Parks, valid publication

## Abstract

Soil samples collected in the Great Smoky Mountains National Park yielded a *Listeria* isolate that could not be classified to the species level. Whole-genome sequence-based average nucleotide identity BLAST and *in silico* DNA-DNA Hybridization analyses confirmed this isolate to be a novel *Listeria sensu stricto* species with the highest similarity to *L. marthii* (ANI=93.9%, isDDH=55.9%). Additional whole-genome-based analysis using the Genome Taxonomy Database Toolkit, an automated program for classifying bacterial genomes, further supported delineation as a novel *Listeria sensu stricto* species, as this tool failed to assign a species identification but identified *L. marthii* as the closest match. Phenotypic and genotypic characterization results indicate that this species is nonpathogenic. Specifically, the novel *Listeria* species described here is phenotypically (i) non-hemolytic and (ii) negative for phosphatidylinositol-specific phospholipase C activity; the draft genome lacks all virulence genes found in the *Listeria* pathogenicity island 1 (LIPI-1), as well as the internalin genes *inlA* and *inlB*. While the type strain for the new species is phenotypically catalase-negative (an unusual characteristic for *Listeria sensu stricto* species), its genome contained an apparently intact catalase gene (*kat*); hence assessment of this phenotype with future isolates will be important. Rapid species identification systems (*Listeria* API, VITEK 2, VITEK MS) misidentified this novel species as either *L. monocytogenes, L. innocua*, or *L. marthii*. We propose the name *L. swaminathanii*, and the type strain is FSL L7-0020^T^ (=ATCC TSD-239^T^).

**IMPORTANCE:** *L. swaminathanii* is a novel *sensu stricto* species that originated from a US National Park, and its place of origin is ultimately preventing this species from achieving valid status. The US National Park Service restricts strain accessibility and open access is currently a prerequisite for species validation. Essentially the only criteria that was not met for *L. swaminathanii* validation is accessibility of the type strain, therefore nomenclature status should not negate the significance of this discovery. As a novel *sensu stricto* species, *L. swaminathanii* expands the group of species whose presence is associated with an increased risk of an *L. monocytogenes* contamination, and therefore could play an important role in public health. While developers of *Listeria* spp. detection methods historically only included validly publish species in their validation studies, *L. swaminathanii* is unequivocally a *sensu stricto* species and should be included as well.

## INTRODUCTION

As of September 2, 2021, there are 26 validly published *Listeria* species. For 58 years, the *Listeria* genus contained only six species (*L. monocytogenes, L. innocua, L. ivanovii, L. seeligeri, L. welshimeri*, and *L. grayi*) that were described between 1926 and 1984 (1-5). Beginning in 2010 with the identification of *L. marthii* (6) and *L. rocourtiae* (7), this genus saw a rapid expansion with a total of 11 species added between 2010 and 2015; in addition to *L. marthii* and *L. rocourtiae, L. fleischmannii* (8, 9), *L. weihenstephanensis* (10), *L. aquatica* (11), *L. cornellensis* (11), *L. floridensis* (11), *L. grandensis* (11), *L. riparia* (11), *L. booriae* (12), and *L. newyorkensis* (12) were added during this period. The 11 newly classified species considerably changed the taxonomy of the genus, notably 10 of these species lacked characteristics historically expected of *Listeria* [e.g., motility, growth at 4°C (13)]; this expanded diversity led to a subdivision into two clades, designated *sensu stricto* and *sensu lato*, based on relatedness to *L. monocytogenes* (14, 15). The *sensu lato* clade is represented by the species showing a more distant relation to *L. monocytogenes*; this clade contains *L. grayi* as well as 10 of the 11 species described between 2010 and 2015. From 2018 to 2020, the *sensu lato* clade continued to expand with the addition of four novel species [*L. costaricensis* - 2018 (16), *L. goaensis* - 2018 (17), *L. thailandensis* – 2019, (18), and *L. valentina* – 2020 (19)]. Between 2010 and 2020, only one species, *L. marthii*, was added to the *sensu stricto* clade; this clade contains *L. monocytogenes* and those species most similar to *L. monocytogenes* (*L. innocua, L. ivanovii, L. seeligeri, L. welshimeri*, and *L. marthii* as of 2020). By 2020, there were 15 novel species (n=1 *sensu stricto*, n=14 *sensu lato*) added to the genus bringing the total number of validly published species to 21.

As part of a project to characterize the prevalence of *Listeria* in soil throughout the contiguous United States, (20), we subsequently identified six novel *Listeria* species (n=4 *sensu stricto*, n=2 *sensu lato*). As of May 2021, only five of these species (*L. cossartiae, L. farberi, L. immobilis, L. portnoyi, L. rustica*) met the criteria to obtain valid standing in the nomenclature (21) according to rules set forth by the International Code of Nomenclature of Prokaryotes [ICNP; (22)], which is governed by the International Committee on Systematics of Prokaryotes [ICSP; (23)]. The sixth species, a novel *sensu stricto*, described here and given the name *L. swaminathanii*, could not be validated. *L. swaminathanii* originated from soil collected in the Great Smoky Mountains National Park (GSMNP) and is therefore the property of the National Park Service by law and subject to the US National Park’s restrictions on discoveries; these restrictions do not allow one to meet ICNP’s requirements for strain accessibility. The five other novel species did not originate from a US National Park and therefore were not subject to US National Park Service (NPS) restrictions. Briefly, for valid standing, ICNP Rule 30 (22) requires that a novel species type strain be deposited into two culture collections where they are made available without restriction. During an October 2021 ICSP executive meeting, the Material Transfer Agreement (MTA) for US National Park isolates was deemed too restrictive (23)and therefore violates ICNP rule 30 (4) (22), which states that type strains cannot be restricted. Hence, valid publication of a novel species solely represented by isolates obtained within a US National Park is currently not possible. A novel species not achieving valid status because it did not meet ICNP requirements (e.g., not obtaining two recognized culture collection certificates) is not uncommon. A 2018 survey of novel species publications (24) found that approximately 150 prokaryotic taxa are identified per year do not have valid standing; this survey calls on authors and journal editors to elevate the status of these species, which would require retroactive adherence to all ICNP rules, and to also limit future addition of non-validly published species (24). Unfortunately, as of November 2021, there is no pathway to valid publication for novel species isolated in US National Parks as neither organization, the US National Park System, nor the ICSP, agreed to change their MTA requirements. At the time of this writing, researchers from the University of Tennessee studying the biodiversity of *Listeria* in the GSMNP also isolated multiple *Listeria* species (n=5) including two strains that could not be classified to the species level (25). Together, these two sampling events suggest the GSMNP could be an invaluable resource for studying *Listeria* diversity; however, current policies that prevent valid publication may deter future research. Novel species isolated in India face similar challenges as the Indian government also imposes restricted access to cultures (26), hence preventing valid publication. Thus, rules intended to protect the rights of discoveries are, in some cases creating a barrier for valid publication of novel species, as place of origin and not scientific evidence may determine if a species obtains official standing in the nomenclature.

## RESULTS

### A soil sample from the Great Smoky Mountains National Park yielded *Listeria* isolates that could not be confirmed to the species level

The novel species described here was isolated from soil collected in the Great Smoky Mountains National Park in North Carolina, USA (Latitude 35.4726543, Longitude −83.851303). A total of 31 *Listeria*-like colonies were isolated that together yielded six different *sigB* allelic types (AT) representing three previously described species including (i) *L. monocytogenes* (1 AT), (ii) *L. innocua* (1 AT), and (iii) *L. booriae* (3 ATs) along with one isolate that could not be classified to the species level (1 AT). The putative novel species is represented by five colonies that all generated the same, novel *sigB* AT (AT 166); these five isolates were designated FSL L7-0020^T^, FSL L7-0021, FSL L7-022, FSL L7-0023, and FSL L7-0024. The observation that the *sigB* AT for these five isolates differed by 8 SNPs from the most closely related *sigB* AT (*L. marthii* AT 42), suggested that these isolates may represent a novel species.

### Whole-genome sequence-based phylogenetic analyses established *L. swaminathanii* is a novel *Listeria sensu stricto* species

To determine whether the five isolates with *sigB* AT 166 represented a novel species, isolate FSL L7-0020^T^ was designated as the type strain with the proposed name *L. swaminathanii* and selected for whole-genome sequencing (WGS) followed by whole-genome-based species delineation assessment via (i) average nucleotide identity using BLAST (ANIb) (27), (ii) *in silico* DNA-DNA Hybridization (isDDH) (28), and (iii) the Genome Taxonomy Database Toolkit (GTDB-Tk) (29-31). The draft genome for *L. swaminathanii* FSL L7-0020^T^ represented 13 contigs and had an N50 length of 1,428,095 bp, an average coverage of 127x, a total length of 2.8 Mb, and G+C content of 38.6 mol%. The total length and G+C content are consistent with the range for current *Listeria sensu stricto* species genomes (2.8 to 3.2 Mb and 34.6 to 41.6 mol%, respectively)(13, 14). The parameters of this draft genome all met the recommended values for taxonomic evaluation set forth by Chun et al. (32).

WGS-based ANIb analysis revealed that *L. swaminathanii* FSL L7-0020^T^ clustered with the *Listeria sensu stricto* clade and showed the highest similarity to *L. marthii* with an ANI value of 93.9% (Fig. 1), which is below the 95% cut-off for species delineation (33). Analysis by WGS-based isDDH also yielded a value below the cut-off for species delineation (<70%) (33). Specifically, *L. swaminathanii* and the most similar reference genome (*L. marthii* FSL S4-120^T^) yielded an isDDH value of 55.9% (confidence interval 53.1-58.6%). Additionally, GTDB-Tk failed to yield a species classification for the *L. swaminathanii* draft genome but did identify *L. marthii* as the most similar genome; the taxonomy of all 34 reference genomes included in the analysis were correctly identified. The phylogenetic tree inferred from the GTDB-Tk output (Fig. 2) positioned *L. swaminathanii* among the *Listeria sensu stricto* clade where it clusters with *L. marthii* and *L. cossartiae*.

**Fig. 1.**
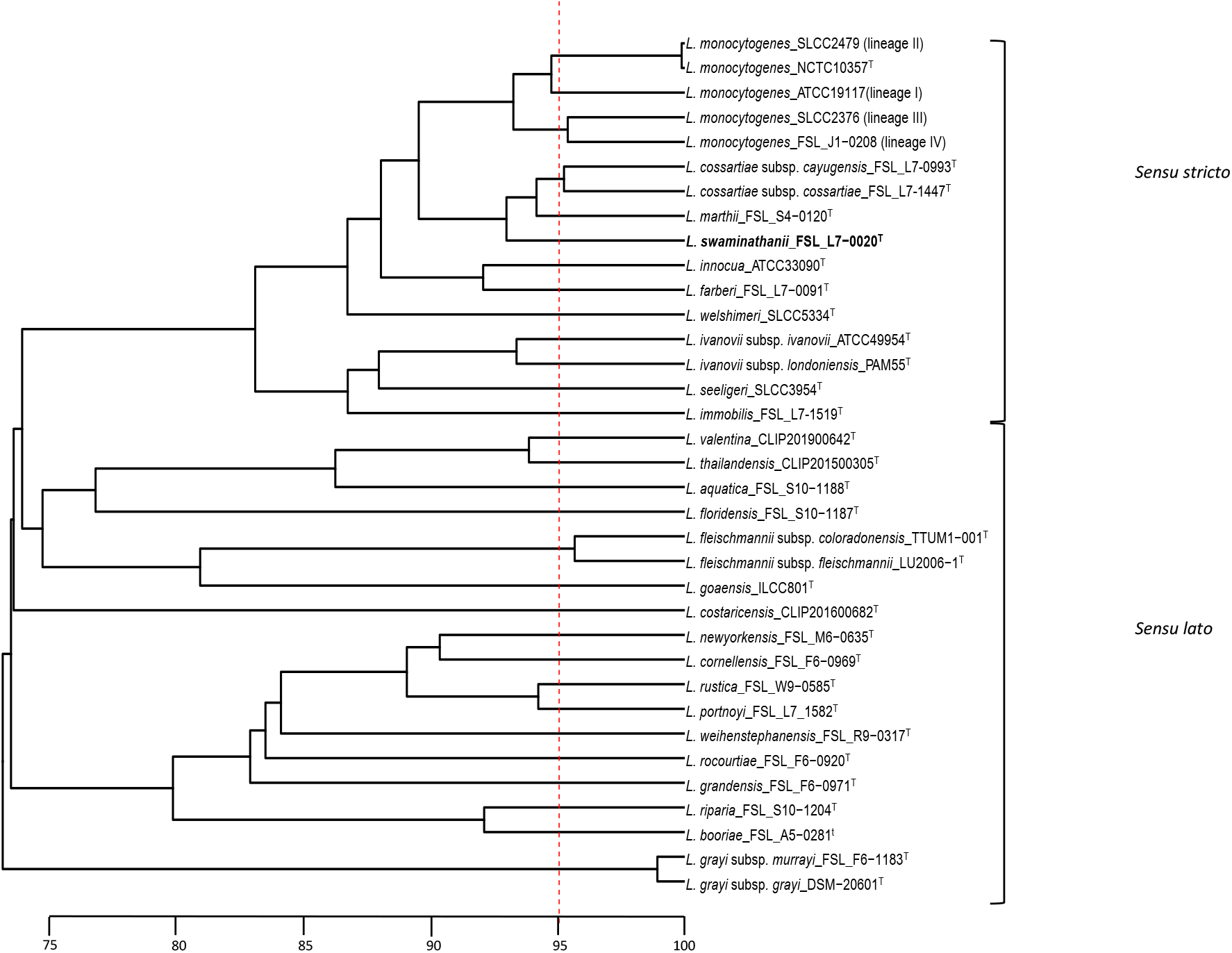
UPGMA dendrogram based on Average Nucleotide Identity BLAST (ANIb) analysis of 34 reference genomes (consisting of the 30 *Listeria* species and subspecies type strains described as of June 11, 2021, and one genome representing each of the four *L. monocytogenes* lineages) and the *L. swaminathanii* FSL L7-0020^T^ draft genome. The vertical red dotted line is placed at 95%, representing the species cut-off. The horizontal scale bar indicates ANI percentage similarity.

**Fig. 2.**
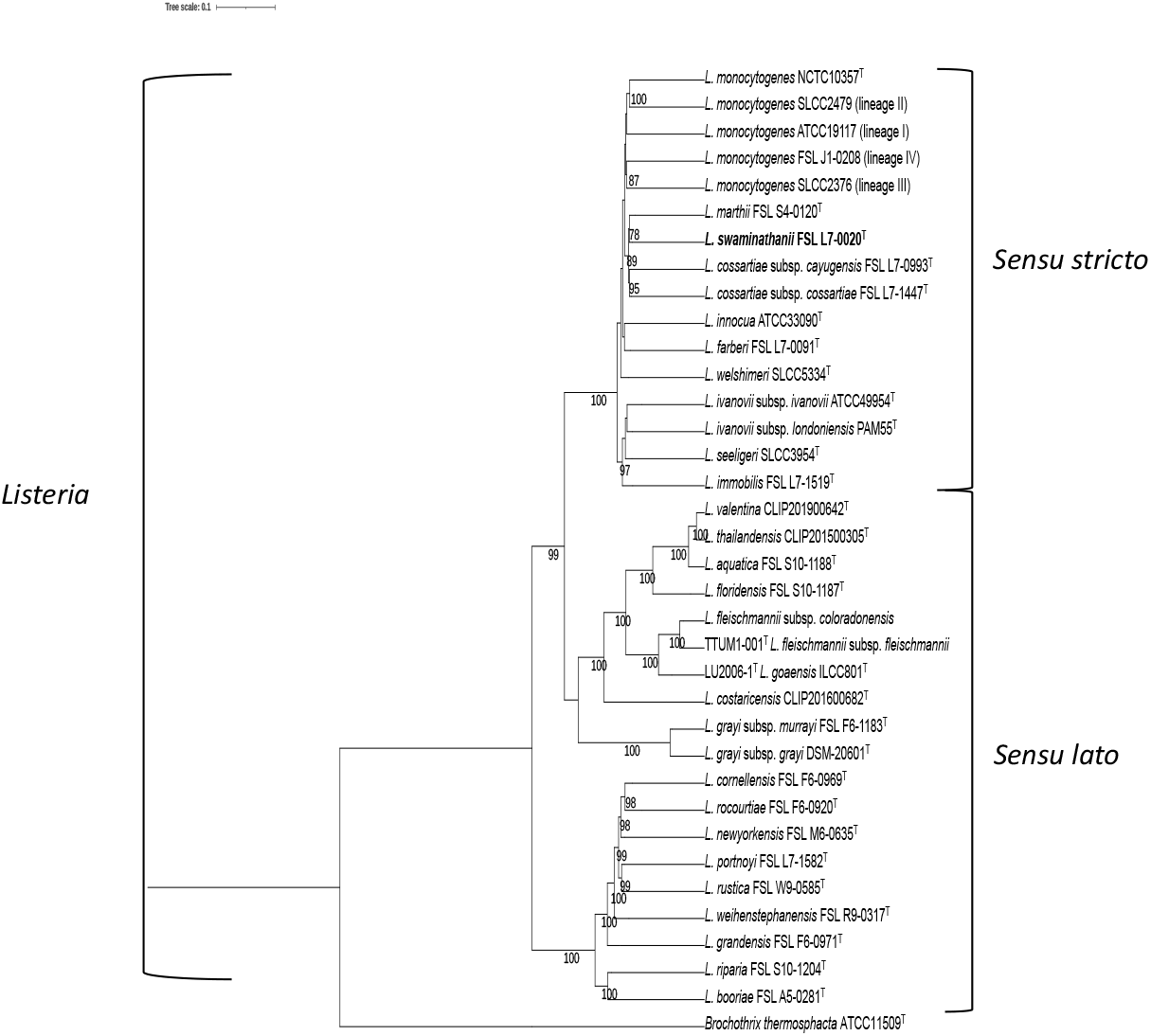
Maximum Likelihood (ML) phylogenetic tree based on the GTDB-Tk analysis of 120 concatenated protein amino acid sequences of the same 34 reference genomes used for ANIb analysis and the *L. swaminathanii* draft genome. The phylogeny was inferred using RAxML v8.2.12 (55), and the best fit model for protein evolution, PROTGAMMAILGF, was determined using ProtTest 3.4.2 (63). The values mapped the nodes represent bootstrap values based on 1,000 replicates; values <70% are not shown. The tree is rooted at the midpoint and includes the outgroup *Brochothrix thermosphacta* ATCC 11509^T^.

### *L. swaminathanii* yielded colony morphologies typical of nonpathogenic *Listeria* spp

Following streaking of an overnight BHI broth culture onto MOX and LMCPM agars, *L. swaminathanii* yielded colonies typical of *Listeria* species (34). When grown on MOX, *L. swaminathanii* FSL L7-0020^T^ yielded black colonies indicative of esculin hydrolysis that were round, had sunken centers, and a black halo; this morphology matches the current description for “typical” *Listeria* spp. growth on MOX (34). Phosphatidylinositol-specific phospholipase C (PI-PLC) activity is a virulence factor presently associated with the pathogenic species *L. monocytogenes* and *L. ivanovii*. On LMCPM agar, PI-PLC activity is detected by the chromogen X-inositol phosphate; colonies positive for PI-PLC activity appear blue-green, negative colonies are white (35). When streaked to LMCPM, *L. swaminathanii* FSL L7-0020^T^ yielded colony morphologies consistent with *Listeria* spp. that are negative for PI-PLC activity. Specifically, *L. swaminathanii* yielded small, round, white colonies on LMCPM. Only *L. monocytogenes* 10403S generated blue-green colonies indicative of PI-PLC activity.

### Except for the catalase negative reaction, *L. swaminathanii* generated the expected biochemical results of a nonpathogenic *Listeria sensu stricto* species

The standard *Listeria* reference method characterization tests we performed included (i) catalase, (ii) oxidase, (iii) Gram staining, (iv) β hemolysis on blood agar, (v) nitrate and nitrite reduction, and (vi) motility. Interestingly, the *L. swaminathanii* type strain FSL L7-0020^T^ was catalase-negative; a characteristic not previously observed with any *sensu stricto* species (6, 13, 21, 34), however several catalase-negative *L. monocytogenes* strains have been reported(36-39). Other than *L. swaminathanii* FSL L7-0020^T^, the only other catalase-negative species reported to date is the recently described *sensu lato* species, *L. costaricensis* (16). The four additional strains representing *L. swaminathanii* (FSL L7-0021, FSL L7-022, FSL L7-0023, and FSL L7-0024) with identical *sigB* ATs were also tested and were also catalase-negative. To further assess the absence of catalase activity, analysis of the draft genome for the *kat* gene was performed, see below for results. Other than the catalase reaction, the oxidase and Gram-stain results were consistent with what is currently expected for *Listeria* spp. (13). Specifically, the *L. swaminathanii* type strain FSL 0020^T^ presented as an oxidase-negative, Gram-positive short rod. Sheep’s Blood Agar (SBA, Becton Dickinson) was used for hemolysis testing. Only *L. monocytogenes* lysed the red blood cells in the agar resulting in a clear zone of β hemolysis (a phenotype associated with *Listeria* pathogenicity); *L. swaminathanii* is non-hemolytic.

None of the *Listeria* species described to date reduce nitrite, while nitrate reduction is currently only observed with the recently described *sensu lato* species (14, 16-19, 21). After the *L. swaminathanii* FSL L7-0020^T^ nitrate broth enrichment was combined with Sulfanilic acid and N, N-Dimethyl-*a*-nathylamine no red color change was observed until the addition of zinc; a red color change was generated when these reagents were combined with the nitrite broth enrichment indicating this species does not reduce nitrate or nitrite. The control strains performed as expected. Specifically, *L. monocytogenes* 10403S did not reduce nitrate or nitrite and *L. booriae* FSL A5-0281^T^ only reduced nitrate.

Motility was assessed both microscopically and following stab inoculation into Motility Test Medium (MTM, Becton Dickinson). For the microscopic method, wet mounts were prepared from BHI agar cultures grown at 25°C and 37°C for 24 h. Motility testing using MTM was performed by stab-inoculating the medium (purchased pre-made in 10 mL screw-capped tubes) with an isolated colony selected from BHI agar followed by incubation at 25°C with observations every 24 h for 7 days. *L. swaminathanii* FSL L7-0020^T^, along with the *L. monocytogenes* positive control, exhibited motility at 25°C with both motility test methods; a tumbling movement was observed microscopically, and an umbrella-like growth pattern was observed following incubation in MTM agar. *L. swaminathanii* FSL L7-0020^T^, along with both control strains, were non-motile at 37°C. To date, *L. costaricensis* is the only *Listeria* species reported to be motile at 37°C (16), and *L. immobilis* is the only *sensu stricto* species the lacks motility at 25°C (21).

### The growth range and optimal growth temperature of *L. swaminathanii* is consistent with that is currently expect of *Listeria* spp

The expected growth range for *Listeria* is currently listed as 0-45°C (13), although exceptions have been identified with several recently described *sensu lato* species that exhibit a narrower temperature range for growth, including eight *sensu lato* species that do not growth at 4°C (14, 16-19) and four species that do not grow at 41°C (7, 10, 21). Presently, all species grow optimally at either 30 or 37°C (14, 16-19, 21). *L. swaminathanii* generated growth at all temperatures tested. The least growth (4.34 log_10_) was recorded after 10 days of incubation at 4°C, and optimal growth was achieved at both 30 and 37°C (9.28 and 9.30 log_10_) after 24 h of incubation. *L. swaminathanii* along with the *L. monocytogenes* 10403S control strain both grew anaerobically. Detailed growth data can be found in Supplementary Table S1.

### *Listeria* API analysis misidentified *L. swaminathanii* as *L*. monocytogenes

*L. swaminathanii* yielded the numeric code 6110 which the apiweb database (bioMérieux V2.0, apiweb version 1.4.0) reported a “very good identification to the genus” with an 80% ID to *L. monocytogenes* and a T value of 0.62. Possible T values range from 0-1.0; the closer the value is to 1.0, the closer the biochemical test results are to what is considered “typical” for the species (40). *L. monocytogenes* was reported as the most likely species due to a negative result for the D-arylamidase activity, referred to as the DIM test (Differentiation of *innocua* and *monocytogenes*) (34). The discordant result leading to a T value of 0.62 is attributed to the negative result for rhamnose fermentation generated by *L. swaminathanii* FSL L7-0020^T^. Differentiation from *L. monocytogenes* may be achieved via both the lack of hemolysis and a catalase-negative tests. The same numeric code (6110) has also been reported for *L. marthii* (6) and *L. cossartiae* subsp. *cossartiae* (21). Phenotypically, *L. swaminathanii* is most easily differentiated from *L. marthii* and *L. cossartiae* subsp. *cossartiae* by the lack of catalase activity. Further differentiating characteristics were determined following the API CH50 analyses described below. *L. monocytogenes* 10403S and *L. innocua* ATCC 33090^T^ were tested to verify *Listeria* API kit performance and generated the expected results for typical strains (numeric codes 6510 and 7510, respectively).

### API 20E results further supports a *Listeria sensu stricto* identification while API CH50 results allow for further differentiation of *L. swaminathanii* from *L. marthii* and *L. cossartiae*

*L. swaminathanii* FSL L7-0020^T^ tested positive for Voges-Proskauer and negative for indole, urease, and H_2_S production via the API 20E test, which is consistent with what is currently expected of *Listeria sensu stricto* species. Specifically, all currently described *sensu stricto* species are Voges-Proskauer negative while all *sensu lato* species are positive. Other than catalase activity, the API CH50 test was needed to phenotypically differentiate *L. swaminathanii* FSL L7-0020^T^ from *L. marthii* and *L. cossartiae*. Specifically, *L. swaminathanii* FSL L7-0020^T^ is negative for fermentation of D-turanose and positive for glycerol and starch utilization while *L. marthii* ferments D-turanose and does not utilize glycerol. The starch result is not commonly used to differentiate *Listeria* and therefore often not reported, however we found *L. swaminathanii* FSL L7-0020^T^’S ability to utilize starch differentiated this isolate from *L. cossartiae* (Tables 1 and S1). A summary of the results commonly reported for *Listeria* are presented in Table 1; additional API CH50 results are provided in Supplementary Table S2.

**Table 1.**
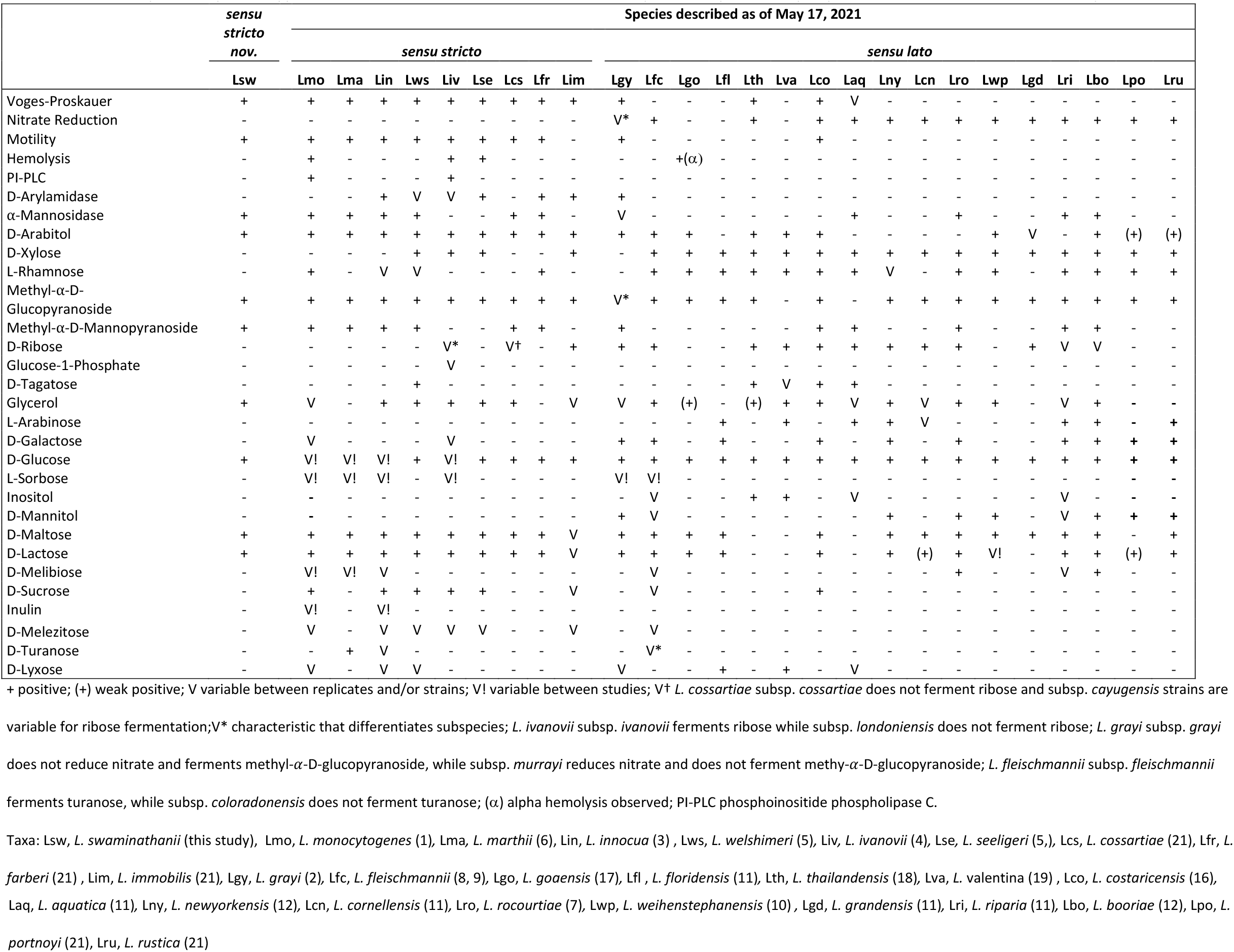
Summary of the phenotypic characteristics of *L. swaminathanii* compared to previously reported characteristics of other species

### VITEK 2 and VITEK MS misidentified the novel *Listeria sensu stricto* species *L. swaminathanii* as well as the other recently reported *L. cossartiae, L. farberi*, and *L. immobilis*

VITEK 2 yielded a good identification for *L. swaminathanii* FSL L7-0020^T^ with a 91% probability of being *L. innocua*. VITEK MS V3.2 identified *L. swaminathanii* L7-0020^T^ as *L. marthii* with a confidence value of 99.9%. Previous publications using VITEK 2 or VITEK MS to characterize novel species did not yield a species identification; however, all these isolates were novel *sensu lato Listeria* species (17, 18). Our data suggest that, unlike novel *sensu lato*, novel *sensu stricto* species could be misidentified given their genetic and phenotypic similarities to the species currently represented in the respective databases. The VITEK 2 database contains strains representing six *Listeria* species (*L. monocytogenes, L. innocua, L. ivanovii, L. seeligeri, L. welshimeri, L. grayi*), and the VITEK MS database contains strains representing seven species, the same set as VITEK 2 plus *L. marthii*. To further evaluate what results are expected for a novel *sensu stricto* species, we also screened the recently described *sensu stricto* species *L. farberi* FSL L7-0091^T^, *L. immobilis* FSL L7-1519^T^, and *L. cossartiae* (both subspecies *cossartiae* FSL L7-1447^T^ and *cayugensis* FSL L7-0993^T^) on both the VITEK 2 and VITEK MS systems. For *L. farberi*, VITEK 2 yielded a low discrimination result with the potential to be *L. innocua, L. monocytogenes*, or *L. welshimeri;* the systems software recommended β-hemolysis, CAMP, and xylose fermentation testing to discriminate the species identification further. *L. farberi* FSL L7-0091^T^ was identified as *L. innocua* with VITEK MS (confidence value 99.9%). *L. immobilis* FSL l7-1519^T^ yielded an excellent identification as *L. ivanovii* with VITEK 2 and was identified as *L. monocytogenes* (confidence value 99.7%) with VITEK MS. *L. cossartiae* subsp. *cossartiae* FSL L7-1447^T^ yielded the same identification reported for *L. swaminathanii* L7-0020^T^ (*L. innocua* with VITEK 2 and *L. marthii* with VITEK MS). *L. cossartiae* subsp. *cayugensis* FSL L7-0993^T^ gave a low discrimination result with VITEK 2 and the possibility of being *L. innocua* or *L. grayi* due to the positive result for ribose fermentation seen with this strain; VITEK MS identified this strain as *L. marthii*.

### Despite absence of catalase activity, the *L. swaminathanii* draft genome contains a full-length *kat* gene

Our initial analyses found that the *L. swaminathanii* draft genome contained a sequence that matched the *L. monocytogenes* partial catalase locus (lmo2785 [*kat*]) from the BIGSdb-*Lm* database with 22 mismatches (over 486 nucleotides). We thus performed a second query using the entire catalase gene sequence (1467 nucleotides; *kat* reference sequence NC_003210.1); the sequence identified by BLASTn was then evaluated using MEGA (41) for premature stop codons. The *L. swaminathanii* draft genome contained a full-length *kat* with no premature stop codons. It hence remains unclear why the type strain (as well as the four closely related isolates obtained from the same sample) are catalase negative; future work on expression of the *kat* gene may be able to resolve this apparent conundrum.

### The *L. swaminathanii* draft genome lacked virulence and sanitizer resistance genes, all flagella genes were detected

The six virulence genes (*prfA, plcA, hly, mpl, actA*, and *plcB*) found on the *Listeria* Pathogenicity Island 1 (LIPI-1) and the internalin genes *inlA* and *inlB* were all absent from the *L. swaminathanii* FSL L7-0020^T^ draft genome. All 26 flagellar genes (Supplementary Table S3) included in the reference database were detected. Genes that were previously reported to confer reduced sensitivity to quaternary ammonium compounds (*qac, bcrABC, ermE*) were not detected.

### *In silico* PCR analysis identified *L. swaminathanii* FSL as a *Listeria* species

A complete *prs* sequence was detected with no mismatches to either the forward or reverse primers supporting *Listeria* species should be detected following PCR analysis. There were no BLAST hits for any of the *L. monocytogenes* serovar specific sequences, indicating that *L. swaminathanii* would not be misidentified as *L. monocytogenes*.

### Description of *Listeria swaminathanii* sp. Nov

*L. swaminathanii* (swa.min.ath.an’i.i NL masc. adj. *swaminathanii* named in honor of Balasubramanian Swaminathan for his contributions to the epidemiology of human listeriosis and laboratory diagnostic methodologies.

*L. swaminathanii* FSL L7-0020^T^ exhibits growth characteristics typical of non-pathogenic *sensu stricto Listeria* spp. except for the catalase reaction, the type strain for this species is catalase-negative. Gram-positive short rods. Oxidase negative. Facultative anaerobe. Presumed to be nonpathogenic based on the absence of hemolysis on SBA, lack of PI-PLC activity on LMCPM, and the absence of six virulence genes (*prfA, plcA, hyl, mpl, actA*, and *plcB)* located on LIPI-1. Colonies on MOX are round, black, approximately 2-3 mm in diameter with a sunken center. Colonies on LMCPM were of similar size and shape as colonies on MOX and are opaque-white in color. Classic umbrella-patterned motility in MTM incubated at 25°C. Tumbling motility is observed microscopically at 25°C. Non-motile at 37°C. Growth occurs between 4-41°C in BHI broth with optimal growth achieved between 30-37°C. Does not reduce nitrate or nitrite. Phenotypically this type strain cannot be differentiated from *L. marthii* or *L. cossartiae* subsp. *cossartiae* using API Listeria (i.e., API numerical profile = 6110) or the biochemical reactions specified in the FDA BAM or ISO 11290-1:2017 methods. Voges-Proskauer positive. *L. swaminathanii* FSL L7-0020^T^ is negative for D-arylamidase activity and positive for α-mannosidase activity. Does not ferment D-xylose, L-rhamnose, D-ribose, glucose-1-phosphate, D-tagatose, L-arabinose, D-galactose, L-sorbose, inositol, D-mannitol, D-melibiose, D-sucrose, inulin, D-melezitose, D-turanose, or D-lyxose. Positive for fermentation of D-arabitol, methyl-a-D-glucopyranoside, methyl-a-D-mannopyranoside, glycerol, D-glucose, D-maltose, D-lactose, and starch. *L. swaminathanii* FSL L7-0020^T^ is differentiated from *L. marthii* by the utilization of glycerol and lack of ability to ferment D-turanose. Differentiation from *L. cossartiae* subsp. *cossartiae* is achieved by the ability to utilize starch.

The draft genome total length is 2.8 Mb with a GC content of 38.7%. Type strain, FSL L7-0020^T^, ATCC TSD-239^T^ was isolated from soil collected in the Great Smoky Mountain National Park, North Carolina, USA, on November 2, 2017. The GenBank/EMBL/DDBJ accession numbers for the 16S rRNA and draft genome sequences for the type strain are MT117895 and JAATOD000000000, respectively.

## DISCUSSION

### Three whole-genome sequence-based classification methods identified *L. swaminathanii* as a novel *Listeria sensu stricto* species, however phenotypic-based differentiation from other species was challenging

*L. swaminathanii* FSL L7-0020^T^ WGS classification analyses confirmed placement of this species within the *Listeria* genus as a novel *sensu stricto* species based on meeting widely accepted species delineation thresholds [ANI <95%, isDDH <70% (33)]. All three WGS-based computational tools (ANIb, isDDH, and GTDB-Tk) used in this study to assess the *L. swaminathanii* FSL L7-0020^T^ draft genome showed this species clusters closest to *L. marthii*. Phenotypically, *L. swaminathanii* FSL L7-0020^T^ may be distinguished from other *sensu stricto* based on the unique catalase-negative attribute; however, this attribute could also lead to a situation where *L. swaminathanii* FSL isolates may not even be identified as *Listeria* as a catalase-positive reaction is often used to confirm *Listeria* to the genus level (34, 42, 43). Interestingly, several cases of human listeriosis have been attributed to catalase-negative *L. monocytogenes* strains (36-39), which supports a need to reduce the reliance on the catalase test for identification of *Listeria* spp. Characterization of additional *L. swaminathanii* isolates will be important to determine whether the catalase negative phenotype is a species-specific phenotype or a strain specific phenotype (similar to what has been reported for *L. monocytogenes*). Beyond the catalase test, phenotypically differentiating *L. swaminathanii* FSL L7-0020^T^ from *L. marthii* and/or *L. cossartiae* subsp. *cossartiae* was difficult; these three species shared the same biochemical results for the species identification tests detailed in the reference methods (β-hemolysis, rhamnose, xylose, mannitol) and generate the same numeric code with API *Listeria* (6110). The API CH50 glycerol, and D-turanose tests provided further species-level discrimination between *L. swaminathanii* FSL L7-0020^T^ and *L. marthii*, and the starch test allowed for further differentiation of *L. swaminathanii* FSL L7-0020^T^ from *L. cossartiae*.

### *L. swaminathanii* along with the other recently described *Listeria sensu stricto* species may not be detected using rapid detection methods and/or commonly used reference methods

For many *Listeria* detection methods (both rapid and cultural), validation studies only included the “classical” six *Listeria* spp. (i.e., *L. monocytogenes, L. innocua, L. ivanovii, L. seeligeri, L. welshimeri*, and *L. grayi*), with some studies also validated with *L. marthii*; it is hence unknown whether many of these assays detect the novel *Listeria* species identified since 2010. This lack of information was less of a concern until recently, given that until 2021 most newly described species (14 out of 15) were classified in the *sensu lato* clade and the food industry is more concerned with detecting *sensu stricto* species as this clade contains *L. monocytogenes* and the species most similar to *L. monocytogenes*. However, with the identification of *L. swaminathanii* and the recent publication of *L. cossartiae, L. farberi*, and *L. immobilis* (21), there are now 10 *Listeria sensu stricto* species, including four that were reported since 2021, which adds urgency to the need to evaluate existing methods for their ability to detect all *Listeria* spp.

Even if a novel *sensu stricto* species is detected by a rapid method (e.g., PCR) or yields *Listeria*-like colonies with cultural methods on the selective and differential agars, there is a strong potential for either misidentification or a false negative with the subsequent confirmatory tests. Currently, catalase and motility tests are utilized to confirm *Listeria* to the genus-level as catalase-positive and motility are considered universal traits to all *sensu stricto* species (13, 34, 42, 43), however there is now sufficient evidence to warrant revising this claim. In addition to the potential for a catalase-negative *Listeria* with *L. swaminathanii* and some strains of *L. moncytogenes* as detailed above, the recently described *sensu stricto* species, *L. immobilis*, is non-motile (21). After the genus-level confirmatory tests, the species-level tests also showed potential for a false negative or misidentification. Presently, the reference methods do not list the expected results for the classic biochemical identification test (β-hemolysis, rhamnose, xylose, and mannitol) for the recently described *sensu stricto* species. As an example, *L. swaminathanii* FSL L7-0020^T^ is non-hemolytic, and negative for rhamnose, xylose, and mannitol fermentation, which is a profile not currently associated with any species in the commonly used reference methods (e.g., FDA BAM, Health Canada, ISO).

### Rapid identification methods showed a strong potential to misidentify the recently described *Listeria sensu stricto species*

As rapid bacterial identification methods are becoming increasingly popular in the food industry, our data highlight the importance of updating reference databases to include at least all the *sensu stricto* species. Unlike the novel *sensu lato*, which historically do not generate acceptable species identifications with the rapid identification methods, the five recently described novel *sensu stricto* species *(L. cossartiae, L. farberi, L. immobilis, L. marthii*) along with the species reported here (*L. swaminathanii*) were all misidentified with the rapid identification methods used in this study. Notably, we saw the potential for a nonpathogenic novel *sensu stricto* to be identified as the pathogenic strains *L. monocytogenes* or *L. ivanovii*; at minimum, this could cause confusion and delays, and worse-case lead to unnecessary product disposals or recalls.

### *L. swaminathanii* should be recognized as a *Listeria* species despite not being able to achieve valid status

In conclusion, while *L. swaminathanii* may not become a validly published species due to restrictions associated with its isolation from a US National Park, its designation as a novel *sensu stricto* species is firmly supported by the results described here. Incorporating this species, along with other recently described species, in *Listeria* method inclusivity studies will be essential to ensure accurate detection and *Listeria* species identification, and consequently better data on the true prevalence of these species. Numerous studies (44-47) have associated the presence of a *Listeria sensu stricto* species to an increased potential for *L. monocytogenes* contamination, hence *L. swaminathanii* may play a key role in public health as detection and subsequent eradication may eliminate a food safety hazard.

## MATERIALS AND METHODS

### *Listeria* isolation and initial identification

As part of a previously reported study evaluating the prevalence of *Listeria* in soil (20), a total of five soil samples were collected from the Great Smoky Mountains National Park and 25g aliquots of each sample were enriched in Buffered *Listeria* Enrichment Broth (BLEB, Becton Dickinson, Frankland Lake, NJ, USA); *Listeria* spp. isolation was conducted as described in the US Food and Drug Administration’s *Bacteriological Analytical Manual* (FDA BAM) Chapter 10 method (34) with one modification: Modified Oxford Agar (MOX, Becton Dickinson) was incubated at 30°C instead of 35°C. From the options for selective and differential chromogenic agars detailed in FDA BAM, we used R&F *Listeria monocytogenes* Chromogenic plating medium (LMCPM, R&F Laboratories, Downers Grove, IL, USA). Following streaking of the five BLEB-enriched soil samples, the MOX and LMCPM agar plates were incubated for 48 h, *Listeria*-like colonies were selected from both plate types and isolated onto Brain Heart Infusion (BHI, Beckton Dickinson) agar. Following isolation onto BHI, species identification was performed using a previously described protocol for PCR amplification and sequencing of the partial *sigB* gene (48).

### Whole-genome sequencing

Genomic DNA was prepared and sequenced, using Illumina’s MiSeq platform, as described in our previous publication (21). The raw sequencing data was assembled, and draft genome quality was assessed using the protocols described by Kovac et al. (49). Briefly, adapter sequences were trimmed using Trimmomatic 0.39 (50), and paired-end reads were assembled *de novo* using SPAdes v3.13.1 (51) with k-mer sizes of 33, 55, 77, 99, 127. Contigs <500 bp were removed, and assembly quality was checked using QUAST v5.0 (52), followed by screening for contamination using Kraken(53).

### Whole-genome-based phylogenic analysis

Whole-genome sequence-based ANIb analysis was conducted on the *L. swaminathanii* FSL L7-0020^T^ draft genome and a set of 34 reference genomes consisting of (i) the 30 type strains of all *Listeria* species and subspecies described as of May 17, 2021, and (ii) a representative for each of the four *L. monocytogenes* lineages (Fig. 1). Pyani (27) was used to calculate pairwise ANI values, and a dendrogram was constructed using the dendextend R package (54). Further analysis by whole-genome sequence-based isDDH was also performed using the Genome-to-Genome Distance Calculator 2.1, formula 2 [identities/ high-scoring segment pair (HSP)] (28). A newer WGS-based computational tool for classifying bacterial genomes, GTDB-Tk [released 2019, (31)], which is a software toolkit that classifies genomes using the Genome Taxonomy Database (GTDB; released in 2018 and updated biannually) (29), was also employed. GTDB infers phylogeny from a set of marker genes made up of 120 bacterial protein genes (bac120) (29), and GTDB-Tk assigns classification based on ANI (calculated with FastANI) and Relative Evolutionary Divergence (RED) scores (29). The same reference genomes used for ANIb were used for the GTDB-Tk analysis of *L. swaminathanii* FSL L7-0020^T^. A phylogenetic tree was inferred from the GTDB-Tk output using RAxML (55), which utilized the alignment of the bac120 protein marker genes from all genomes assessed (the 34 reference genomes and the *L. swaminathanii* draft genome) along with *Brochothrix thermosphacta* ATCC 11509^T^ (output group). The tree was visualized using Figtree (56).

### Phenotypic analyses

Phenotypic characterizations of *L. swaminathanii* FSL L7-0020^T^ were carried out using BHI agar cultures streaked from a frozen stock culture (stored at −80°C in BHI broth supplemented with 15% glycerol), followed by incubation at 30°C for 24-36 h. The tests performed include the species classification tests outlined in commonly used reference methods for *Listeria* species identification [the FDA BAM Chapter 10 (34)and ISO 11290:2017 (43)] beginning with colony morphology observations on selective and differential agar. Specifically, colony morphologies were assessed by streaking an overnight BHI broth culture onto MOX and LMCPM agars, followed by incubation at 35°C for 48 h. *L. monocytogenes* 10403S and *L. innocua* ATCC 33090^T^, were included as positive and negative controls respectively. Following morphology assessments, the reference method characterization tests we performed included (i) catalase, (ii) oxidase, (iii) Gram staining, (iv) β hemolysis on blood agar, (v) nitrate and nitrite reduction, and (vi) motility. Two biological replicates were performed for each test. Catalase, oxidase, Gram-staining, and β hemolysis analyses were conducted as described in the reference methods (34, 43) using colonies grown on BHI agar as described above. *L. monocytogenes* 10403S and *L. booriae* FSL A5-0281^T^ were included as negative and positive controls, respectively.

Nitrate and nitrite reduction tests were performed in parallel using a method described by Buxton et al. (57). Briefly, a heavy inoculum from a freshly prepared BHI agar culture was inoculated into both Nitrite and Nitrate broths [prepared according to Buxton et al. (57)], followed by incubation at 35°C. Analyses were performed after 24 h and again after five days of incubation. Following incubation, aliquots of each culture were separately added to commercially prepared reagents of Sulfanilic acid and N, N-Dimethyl-*a*-nathylamine (commercially named NIT1 and NIT2, respectively, bioMérieux). When combined with NIT1 and NIT2, a red color change indicates the presence of nitrite in the nitrate enrichment broth (indicating that nitrate was reduced) or in the nitrite enrichment broth (indicating that nitrite was not reduced). Powdered zinc (bioMérieux), which reduces nitrate to nitrite, was added to the nitrate enrichments that did not exhibit a red color change. Following the addition of zinc, a red color change indicates nitrate was present; no color change indicates nitrate has been completely reduced to, nitric oxide, nitrous oxide, or molecular nitrogen (i.e., the species reduced nitrate).

Motility was assessed both microscopically and following stab inoculation into Motility Test Medium (MTM, Becton Dickinson). For the microscopic method, wet mounts were prepared from BHI agar cultures grown at 25°C and 37°C for 24 h. Motility testing using MTM was performed by stab-inoculating the medium (purchased pre-made in 10 mL screw-capped tubes) with an isolated colony selected from BHI agar followed by incubation at 25°C with observations every 24 h for 7 days.

### Growth experiments

We assessed growth of *L. swaminathanii* L7-0020^T^ at 4, 22, 30, 37, and 41°C by inoculating BHI broth with 30-300 CFU/mL, followed by incubation at the specified temperatures without shaking. The inoculum was verified by spread plating onto BHI agar followed by incubation for 24-36 h at 30°C. The BHI cultures incubated at 4°C were enumerated after 10 and 14 days, BHI cultures incubated at all other temperatures were enumerated after 24 and 48 h of incubation. *L. monocytogenes* 10403S was included as a positive control. Enumerations were carried out by serial diluting and spread plating 100 µL in duplicate onto BHI agar, followed by incubation at 30°C for 24-36 h. After incubation, colonies on BHI agar were counted using the automated SphereFlash colony counter (IUL Micro, Barcelona, Spain). Relative growth for each temperature was calculated as the average of the duplicate counts minus the starting inoculum. Anaerobic growth was assessed by streaking to BHI agar followed by incubation at 30°C for 24h under anaerobic conditions. The growth experiments were performed in two biological replicates.

### API *Listeria*, CH50, and 20E test kit analyses

The API kit tests were performed per the manufacturer’s instruction. Specifically, the API *Listeria* strips were prepared and incubated at 35°C for 18-24h. For API CH50, *L. swaminathanii* L7-0020^T^ was suspended in CHB/E medium, and the strips were inoculated per the manufacture’s instruction, followed by aerobic incubation at 30°C and assessment of results at both 24 and 48 h (reactions that were positive at either time point were considered positive). The API 20E was utilized because it includes tests classically used to characterize *Listeria* spp. to the genus level, including (i) Voges-Proskauer, (ii) indole, (iii) urease, and (iv) H_2_S production. For API 20E, *L. swaminathanii* was suspended in NaCl 0.5% Medium (bioMérieux), and the strip was inoculated per the manufacturer’s instruction, followed by incubation at 35°C for 24 h. API testing was performed in two biological replicates.

### VITEK 2 and VITEK MS analyses

*L. swaminathanii* FSL L7-0020^T^ along with the type strains for the recently reported novel *sensu stricto* species *L. cossartiae* (susp. *cossartiae* FSL L7-1447^T^ and subsp. *cayugensis* FSL L7-0993^T^), *L. farberi* FSL L7-0091^T^, and *L. immobilis* FSL L7-1519^T^ were prepared and processed on the VITEK 2 (bioMérieux) V7.01 and VITEK MS (bioMérieux) V3.2 automated identification systems per the manufacturer’s instructions. After inoculating the GP (Gram-Positive) reagent card, the VITEK-2 system automatically assesses 64 biochemical reactions that are compared to a database to generate an identification (58). VITEK MS is a Matrix-Assisted Laser Desorption Ionization Time-Of-Flight (MALDI-TOF) mass spectrometry method that automatically compares an isolates spectrum to a database (59). For VITEK 2, unknown biochemical patterns are reported as outside the scope of the database. For VITEK MS, the resulting spectra are assigned a percent probability ranging from 60 to 99.9%. Values <%60 are assigned to spectra too different from any in the database, such that no possible identification is provided.

### *Listeria* catalase, virulence, flagellar, and sanitizer resistance gene analyses

The nucleotide sequences for catalase, virulence, flagella, and sanitizer resistance genes for BLASTn reference databases were downloaded from the open-access Institut Pasteur database, BIGSdb-*Lm*, as described by Moura et al. (60) and Ragon et al. (61). Specifically, from BIGSdb-*Lm*, we obtained a partial catalase sequence from the MLST scheme, the flagellar sequences from the cgMLST1748 scheme, the virulence gene sequences from the Virulence scheme, and the sanitizer resistance sequences from the Metal and Disinfectant Resistance scheme.

### In silico PCR for Listeria monocytogenes serovars

*In silico* PCR (isPCR) using the primers described by Doumith et al. (62) for differentiation of *L. monocytogenes* serovars was also performed. We first performed a BLASTn query of the *L. swaminathanii* draft genome against the *L. monocytogenes* serovar specific genes (*lmo0737, lmo118*, ORF2819, ORF2110) and a gene common to all described species (*prs*). The reference sequences were obtained from the BIGSdb-*Lm* database described above from the PCR Serogroup scheme. The sequences obtained from the query were subsequently tested against the primer sequences for an isPCR analysis.

